# Aligning gene trees with family trees

**DOI:** 10.64898/2025.12.11.693727

**Authors:** Andrii Serdiuk, Claudia Moreau, Jean Mathieu, Élise Duchesne, Cynthia Gagnon, Simon Girard, Simon Gravel

## Abstract

Relatedness between individuals can be measured at the genealogical level (by describing shared ancestors in a pedigree) or at a genetic level (by describing shared haplotypes across the genome). The shared haplotype structure can be conveniently summarized as a sequence of trees along the genome, where each tree describes the last shared genetic ancestors between individuals at a locus. While many tools exist to infer tree sequences from genetic data, and many large pedigree datasets are available, few tools exist to identify the relationship between the two – finding which genetic ancestor corresponds to which pedigree ancestor, and conversely.

In this paper, we propose an algorithm to solve this problem by providing, for each genetic tree, the list of all consistent ancestry paths within a genealogical tree. We also provide variants of the algorithm with moderate robustness to both tree sequence and pedigree errors. We demonstrate the scalability of our approach on the BALSAC genealogical dataset, which includes millions of individuals in Quebec, Canada. We find that 20 carriers usually provide enough information to reliably identify a common ancestor 15 generations ago or find the parent of origin of alleles among probands, but the number of possible ancestry paths within the pedigree can remain large. We apply the method to reconstruct the inheritance of a causal allele for type 1 myotonic dystrophy.

## Introduction

Given a set of carriers of a genetic allele and their pedigree, we are interested in figuring out the inheritance path of the allele within the pedigree. Versions of this problem have occupied the minds of young parents and classical thinkers trying to figure out whom babies most resemble. In the age of genetics, familial linkage studies have sought to infer transmission patterns within pedigrees. For example, MORGAN (Wijsman et al. 2006) infers the transmission history within the pedigree by using a Markov Chain Monte Carlo (MCMC) model.

Here we are interested in scaling such approaches to pedigrees including millions of individuals, as we may find in specific human populations (e.g., in Quebec or Iceland), and in many agricultural populations. For concreteness, we will focus in this work on the population of Quebec, where the BALSAC project (Vézina and Bournival 2020) has compiled a pedigree linking approximately 5 million individuals to about 6500 genealogical founders, primarily French settlers to New France. Solving this can be helpful to understand the evolution of disease variants and establish their regional prevalence (Nelson *et al*. 2018; Mejia-Garcia *et al*. 2025).

Classical pedigree approaches do not scale to population-scale pedigrees. One scalable approach is ISGen (Nelson *et al*. 2018), which uses importance sampling to identify ancestry paths consistent with a set of known carriers and has been used successfully on a variety of rare variants (Mejia-Garcia *et al*. 2025; Chong *et al*. 2025). However, ISGen does not take into account the structure of shared haplotypes among carriers. As a result, it has to integrate over a very large number of possible ancestral paths, reducing accuracy and making it computationally intensive.

The haplotype structure is encoded in the ancestral recombination graph (ARG). Recent progress in ARG reconstruction from genetic data (Zhang *et al*. 2023a; Kelleher *et al*. 2019; Speidel *et al*. 2019; Palamara *et al*. 2018) enables population- and genome-scale inference. These ARG inference methods integrate haplotype information from across the genome to predict a gene tree at a specific locus. Vertices in this tree correspond to inferred recent common ancestors between carriers.

Given the true ARG and the correct pedigree, the problem of matching the two can be framed as a graph alignment problem: To each genetic ancestor (a vertex in the ARG), we want to associate a genealogical ancestor (a vertex in the pedigree), or place it outside the pedigree. We found it to be a hard problem. In this work we will focus on the problem of aligning a single gene tree, that is, the ARG restricted to a single locus. We will propose an algorithm that we believe is, in principle, scalable to the entire ARGs.

Concretely, we will use an ARG inferred by an existing ARG-inference strategy, extract the tree at a relevant locus, consider a clade of that tree inferred to share a recent common ancestor, and align that clade to a pedigree. Even though this clade only describes ancestry at a given locus, it is inferred using the haplotype structure – e.g., siblings share on average a longer haplotype than distant cousins and are expected to be placed more often near each other in the tree.

In simulations, even with perfect data, we often find many possible alignments due to the intertwined familial relationships in a centuries-deep pedigree. We will therefore seek an alignment strategy that lists all possible compatible alignments, which enables Bayesian integration over alignment uncertainty. Interestingly, we find that this multiplicity of solutions ensures a degree of robustness against errors in genealogical records and ARG inference, and we find in applications to a causal allele for type 1 myotonic dystrophy that the approach accurately predicts the parent-of-origin of carriers even in cases where the inferred common ancestors are distant.

### The ARG and pedigree alignment problem

A pedigree *P* is a graph in which individuals are represented as vertices, and edges connect offspring to their two parents. The haploid pedigree graph represents the same information, but with each individual being represented as two vertices (a maternally-inherited haploid copy of the genome, and a paternally-inherited one). We refer to each haploid pedigree vertex as a *pedigree ploid*.

An ARG is a complex structure representing the history of coalescent and recombination events at every locus. Vertices in the ARG represent shared inheritance from an ancestral individual or, more precisely, from an ancestral pedigree ploid. Edges in the ARG represent inheritance from a series of ploids. This is most easily visualized by considering a tree obtained by restricting the ARG to a single locus. In Figure 1, the edge 6–7 corresponds to the blue path between the ploids 6 and 7 in the ploid pedigree.

**Figure 1.**
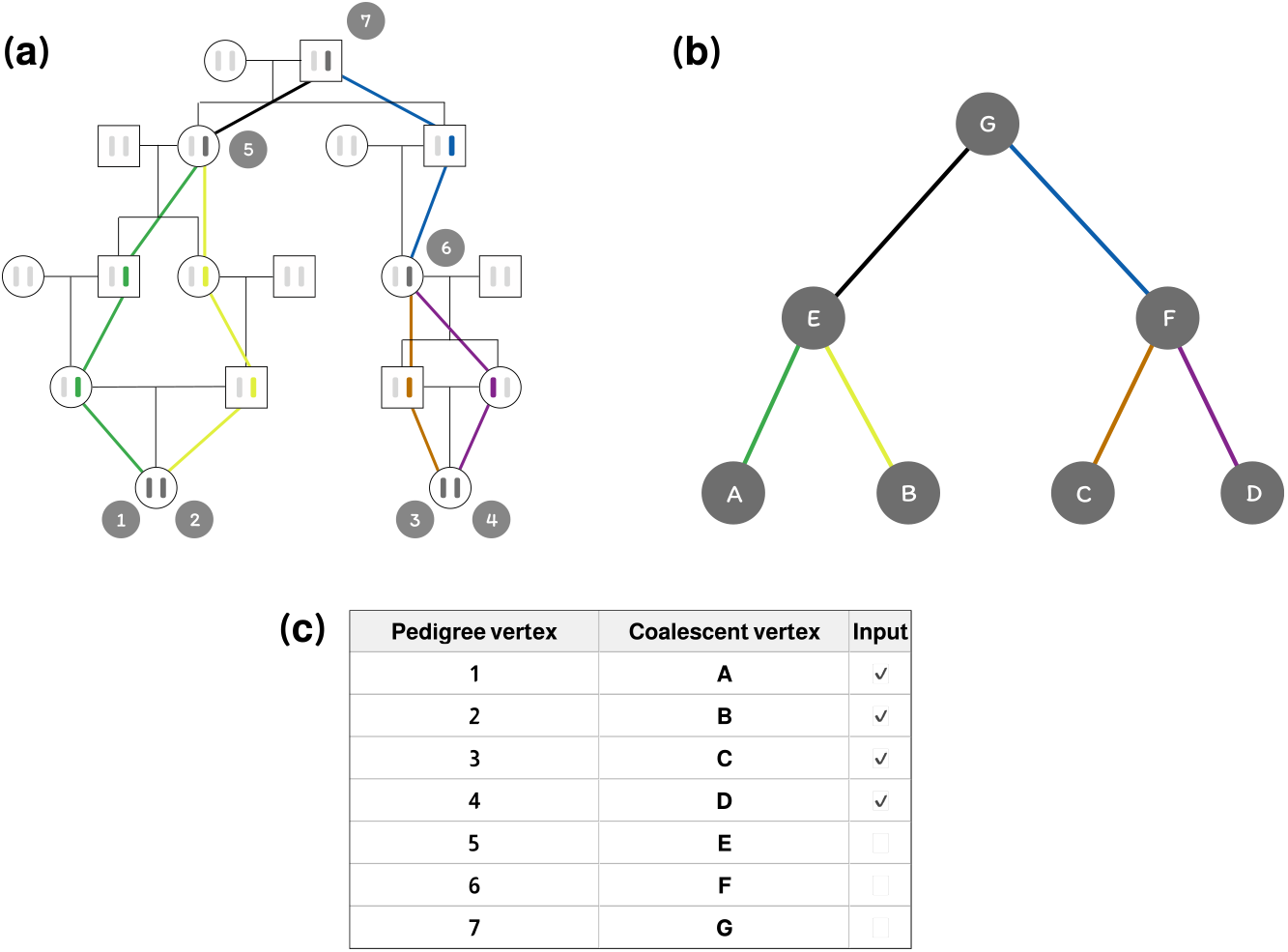
Visualization of the alignment problem, where (a) and (b) represent the input pedigree and the coalescent tree, respectively; (c) shows the input mapping *g* as part of the resulting vertex alignment.

### Problem statement

#### Input

- A pedigree *P*;
- A local tree *T* describing the genetic transmissions in *P*, where *L*_*T*_ is the set of leaf vertices in *T*;
- For each leaf vertex in *L*_*T*_, a set of possible pedigree ploids in *P*: *g* : *L*_*T*_ → 2^*V*(*P*)^ \ {Ø}.

In practice, while studying rare mutations, we almost always know to which individual a specific tree leaf corresponds. However, we often do not know from which parent the individual inherited the mutation. In cases where this is known, we say that we’re working in the *phased regime*, and *g* maps a tree vertex to a pedigree ploid in *P*.

Otherwise, when the parent carrying the mutation is unknown, we say that we’re working in the *unphased regime*, and *g* maps a tree leaf to both pedigree ploids of the same individual.

#### Output

Given the input described above, we want to find the exhaustive list of all the possible histories of genetic transmissions in *P* that respect *T*. Notice that every tree edge represents one series of genetic transmissions between an individual and their MRCA. That is, every history of genetic transmissions in *P* can be thought of as a map assigning every edge in the tree to a pedigree path.

#### Edge alignment

Using the intuition described above, we define the notion of an *edge alignment*. An edge alignment is a map Γ that assigns each edge in *T* to a path in *P* such that the image of Γ represents a consistent history of genetic transmissions (see Appendix A for more details). Therefore, we want to find all such functions to fully solve the problem.

Notice, however, that the space of all possible edge alignments grows rapidly as the size of the input graphs increases. Theoretically, this can make the problem computationally intractable. For this reason, we first consider a coarser but more tractable definition of an alignment.

#### Vertex alignment

Rather than assigning each edge in *T* to a path in *P*, we can assign every vertex in *T* to a vertex in *P*, ignoring details of the path that was followed. We refer to this task as a *vertex alignments*. Notice that one vertex alignment can represent multiple edge alignments, potentially reducing memory requirements. In this context, a vertex alignment is *valid* if there is a valid edge alignment that implies the given vertex alignment. Figure 1 shows an example of a vertex alignment and a corresponding edge alignment.

### Algorithm description

The main idea behind the algorithm is to first find a superset of the valid vertex alignments using a divide-and-conquer approach inspired from Sankoff and Rousseau (Sankoff and Rousseau 1975; Grundler *et al*. 2025; López Sánchez *et al*. 2025). We then find the corresponding edge alignments, filtering out those vertex alignments that don’t have any valid edge alignments. Here and further, let *T*_*v*_ represent the subtree of *T* obtained by cutting *T* above vertex *v*.

The general structure of the algorithms is as follows:

(a) Find a topological ordering of *T*, so that children are always processed before parents;
(b) Climbing the tree *T*, for a vertex *v* and its child vertices *v*_1_, …, *v*_*n*_, find all the potential vertex alignments for the subtree *T*_*v*_ using the solutions for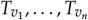;
(c) When the root of *T* is reached, reconstruct all the potential vertex alignments. For every potential vertex alignment, check correctness by validating that there exists at least one edge alignment.

#### Step (a)

Any topological ordering of *T* is suitable, and we define an ordering based on the maximal distance of vertices to the leaves of the tree.

#### Step (b)

Next, we need to find all the possible sub-alignments for *T*_*v*_ given that we know all the possible alignments for the *v*’s children vertices. Given a fixed assignment for each child node, we list individuals who can be a most recent common ancestor to the corresponding individuals, which is required for valid assignments for *v* (see Appendix B).

We solve this problem for each allowed assignment of the child nodes. Since the allowed assignments form the Cartesian product of the allowed assignment for each child, the number of assignments can be large. Fortunately, there are various traversal optimizations that we can use (see Appendix B). The assignments for *v* is, by construction, a common ancestor to all the pedigree individuals assigned to the tree leaves, so that the number of candidates does not grow uncontrollably.

Using this approach, we proceed with the inference until we reach the root of *T*.

#### Step (c)

The procedure in step (b) is guaranteed to include all valid vertex alignments. However, it could also include invalid vertex alignments, i.e., ones that do not have a corresponding edge alignment: the paths from child nodes to parent nodes may collide. To confirm validity, we need to find at least one edge alignment for each vertex alignment. Fortunately, this can be performed efficiently as well using a maximum-flow algorithm (see Appendix A for more details). We can also enumerate all compatible edge alignments if desired, although this can be more computationally demanding (see Discussion).

#### Algorithm scalability

The scalability of the enumeration algorithm depends on how well-constrained the problem is. If the number of constraints is sufficient, we can quickly eliminate the majority of candidates on the first levels of the alignments, leaving us with a small search space. Conversely, if the constraints are insufficient, the search space can rapidly expand, leading to an impractically long search time.

To estimate the potential running time of the algorithm, we used the msprime package (Baumdicker *et al*. 2021; Nelson *et al*. 2020; Anderson-Trocmé *et al*. 2023) to simulate data conditioned on the BALSAC pedigree (Vézina and Bournival 2020) with 3353558 probands. We then selected the largest clade with 30593 vertices (22838 probands) from the simulated coalescent tree and applied the alignment between the clade and the pedigree, assuming known phasing.

The implemented algorithm successfully solved the alignment problem for this input in approximately 11 minutes, identifying 8 distinct vertex alignments with each having only 1 possible edge alignment. In this case, the problem is well-constrained, as the number of resulting alignments is small, and the alignments themselves are very similar. Specifically, the tree’s root vertex could only be assigned to one of the 4 ploids of a founding couple (the natural symmetry of the problem can be observed in Figure 2 where all the green pedigree ploids are consistent MRCAs).

**Figure 2.**
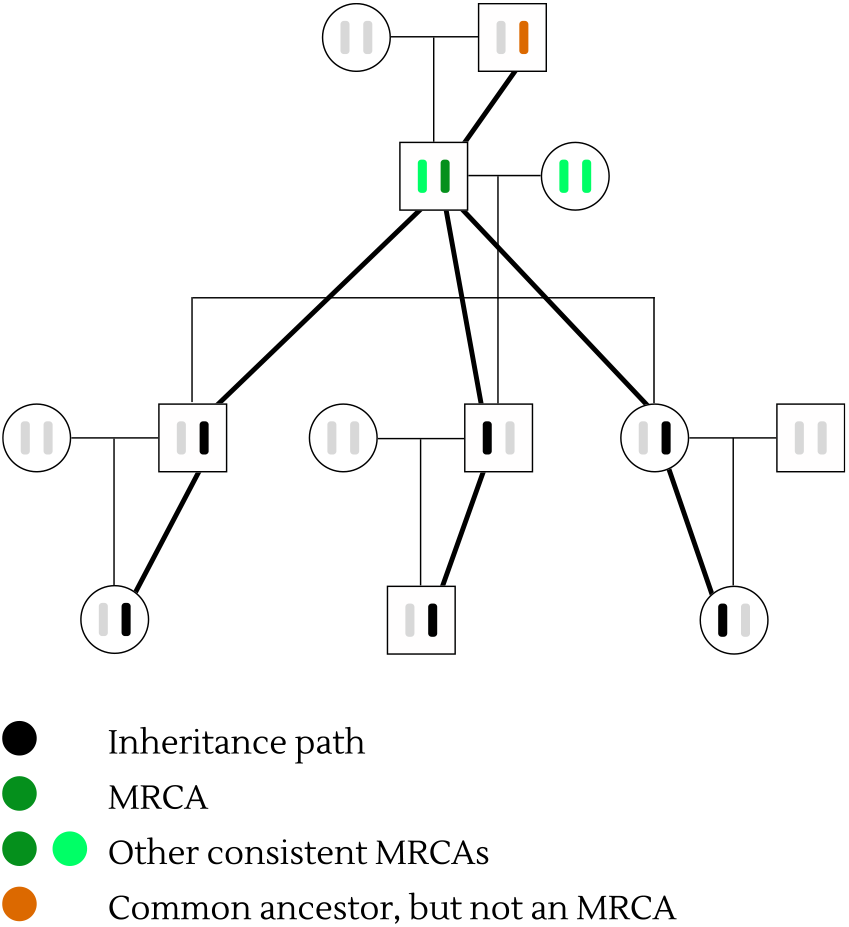
A pedigree MRCA is an individual who can be a coalescent MRCA.

## Results

In this section, we simulate more realistic data and run the algorithm under different conditions to assess the algorithm’s scalability to large graphs and understand how well-constrained the alignment problem is.

### Simulation setup

We consider the participants of the CARTaGENE cohort who have consented to link their genetic and genealogical data(McClelland et al. 2025), extract the ascending genealogies via the BALSAC project (Vézina and Bournival 2020), and use again the msprime package to simulate a coalescent tree conditional on the pedigree (Baumdicker et al. 2021; Nelson *et al*. 2020; Anderson-Trocmé et al. 2023). Then, for each number of probands between 2 and 38, we select up to 10 clades from the simulated tree, resulting in a total of 181 clades. The distribution of the number of clades by the proband number is shown in S3.

### Proband selection experiment

The purpose of this experiment is to quantify how many alignments are found as a function of the number of probands. Specifically, we take the simulated data from the previous section and run the alignment algorithm with the pedigree and every simulated tree in phased and unphased regimes.

Figures 3, S1 illustrate how the number of alignments changes as the number of probands increases under the unphased and phased regimes, respectively.

**Figure 3.**
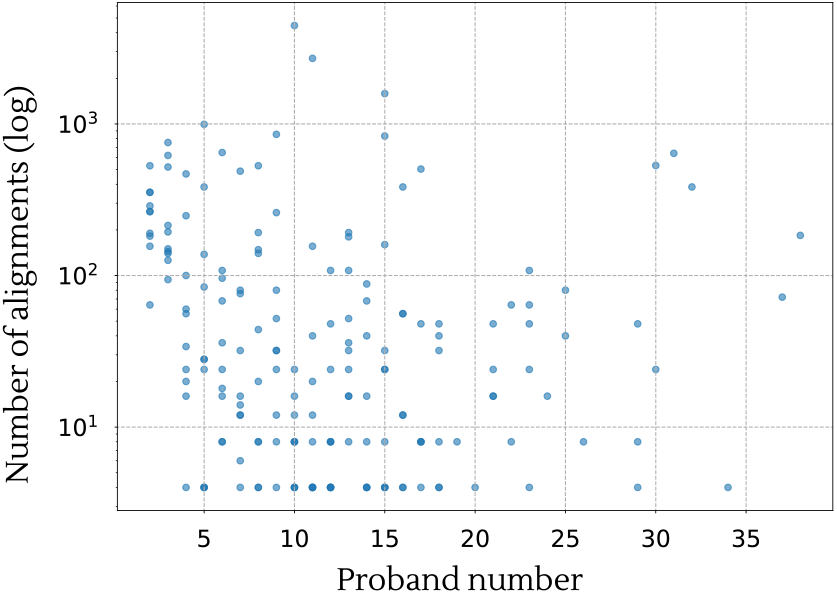
Number of alignments vs proband number (unphased regime).

Interestingly, while the number of valid alignments tends to decrease with an increased number of probands, the number of alignments can remain high even with a large number of probands. We interpret this as resulting from the competing effects of an increased number of constraints offered by additional probands and the increased number of variables to infer: more probands means more edges in the tree, thus more paths to infer. If an additional proband can be connected in multiple ways to existing edge alignments, for example, it can vastly increase the number of possible distinct alignments, even though the average distance between the inferred and true alignments tends to decrease.

### Measuring success

Indeed, even in cases when we have a large number of solutions, the results often share a significant proportion (60-90%) of identical vertex assignments among them. By inspection, we found that this typically happens when a handful of vertices in the gene tree can be frequently reassigned to a different pedigree ploid without altering the topology of the tree. If a few such poorly constrained vertices exist, one can end up with a large number of similar solutions that differ from each other only by the permutation of a handful of poorly constrained vertex assignments. This behavior is typical in large constraint satisfaction problems, where it is common to find a large number of solutions that are clustered around the desired solution even in well-constrained problems (Mézard *et al*. 2005).

We are therefore interested in measuring consistency among alignments, for example, by considering how often they agree on the identity of the last common ancestor to all carriers. Due to the symmetry between the ploids of the true carrier and their spouse (in monogamous unions) (see Figure 2), we expect to find the 4 symmetrical ploids as possible ancestors. We can then inspect how many other individuals tend to be valid assignments to the root, as the clade size increases. Figure 4 shows that, as the proband number becomes larger, we tend to find only the correct couple.

**Figure 4.**
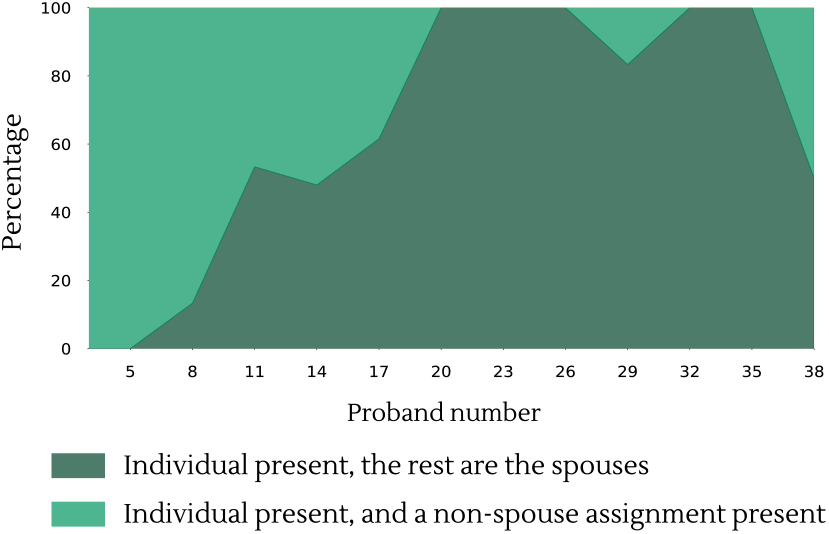
The precision of the root assignment on the data from Simulation setup in the unphased regime.

### A probabilistic model

Given a set of siblings carrying a rare variant, we may expect that the most likely common ancestor is one of the two parents, rather than a shared ancestor 15 generations ago. To formalize this intuition, consider a model in which we observe that a set of *C* probands share a rare variant or a long haplotype block.

Based on some modelling assumptions, we make the hypothesis *X* that an inferred marginal tree has the correct topology for these probands, and that the common ancestor is within the pedigree. We are interested in the posterior probability of a given ancestry path Γ_*C*_ for these individuals, given this hypothesis. Using Bayes’ rule,

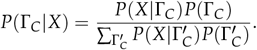

Further, the likelihood *P*(*X* |Γ_*C*_) is exactly one for paths that correspond to valid edge alignments, so that

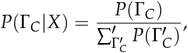

where the sum is taken only on the valid edge alignments.

Under these assumptions, we can thus estimate a posterior probability of a vertex alignment. Let *F* = {*f*_1_, …, *f*_*n*_ } be the set of all vertex alignments between a given coalescent tree and a pedigree. Then, let 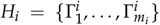 be the set of edge alignments corresponding to a vertex alignment *f*_*i*_. Using the built probability model, the probability of an edge alignment is:

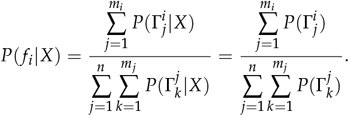

That is, the probability of a vertex alignment is the sum of the corresponding edge alignments’ prior probabilities divided by the sum of all the edge (vertex) alignments’ probabilities.

The prior probability of an ancestral path is

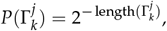

where length(Γ) is the number of meiosis in the edge alignment. Indeed, when conditioning on a pedigree, shorter paths have a higher prior because they require fewer choices.

Similarly, we can define the probability of a pedigree vertex *v* to be a carrier (that is, to be included in an edge alignment). For brevity, let’s use 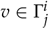 to indicate that *v* is included in the edge alignment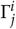

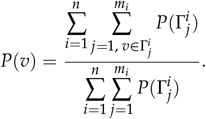

We use this Bayesian model for two purposes. We first validated that, for clades with a high number of vertex alignments, high-posterior solutions are more likely to be near the correct solution (Figure 5). To see how often the posteriors can be used to identify true solutions, we conducted another experiment for the data from Simulation setup.

**Figure 5.**
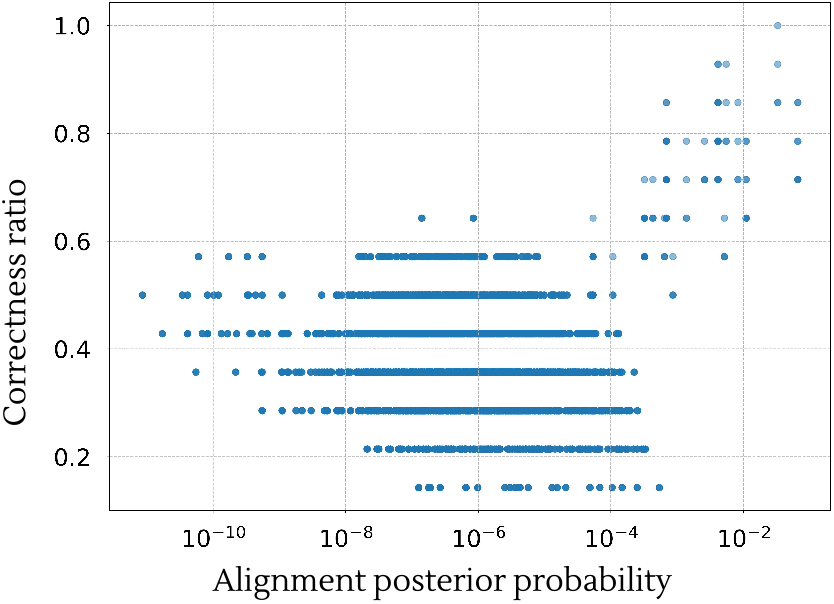
Solutions with high posterior solutions are closer to the true solution for a clade from the Simulation setup with 11312 vertex alignments. Each dot represents a single vertex alignment.

Specifically, we defined a minimal credible set by selecting the smallest set of vertex alignments whose cumulative posterior is above a threshold *β*, typically 95% (see Figure 6). We find that the posterior is usually informative, but in most cases it only mildly reduces the size of the confidence set. In a few cases, however, the posterior is extremely informative (especially when the number of alignments is large). Our interpretation is that many clades in a founder population like the one we consider here are star-like. Thus, many of the possible alignments are also star-like, meaning that most alignments will have comparable lengths. By contrast, in cases where we have non-starlike alignment, the posterior can be very informative.

**Figure 6.**
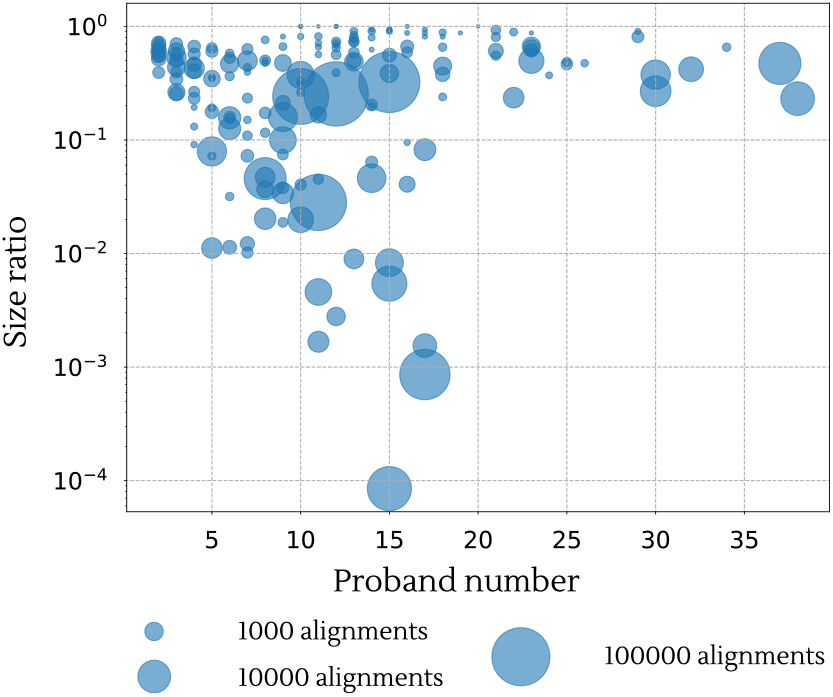
Credible set experiment for the data from Simulation setup. To measure how informative the posterior calculation is, we show for each clade alignment the relative size of the 95% credible sets to the size of credible sets under a uniform posterior. Lower values represent more informative posteriors. The size of the dot is proportional to the number of vertex alignments for that clade.

Second, to compute the expectation of maternal and paternal inheritance for the probands (i.e., parent-of-origin) for the same data from Simulation setup (Figures 7 and S2). Because posterior probability calculation requires the computation of all edge alignments, the computation can be slow for very poorly constrained problems. In Figures 7 and S2, two clades with 16 and 31 probands had an enumeration time of more than a week, and were not included (see Occasional long inference time for edge alignments).

**Figure 7.**
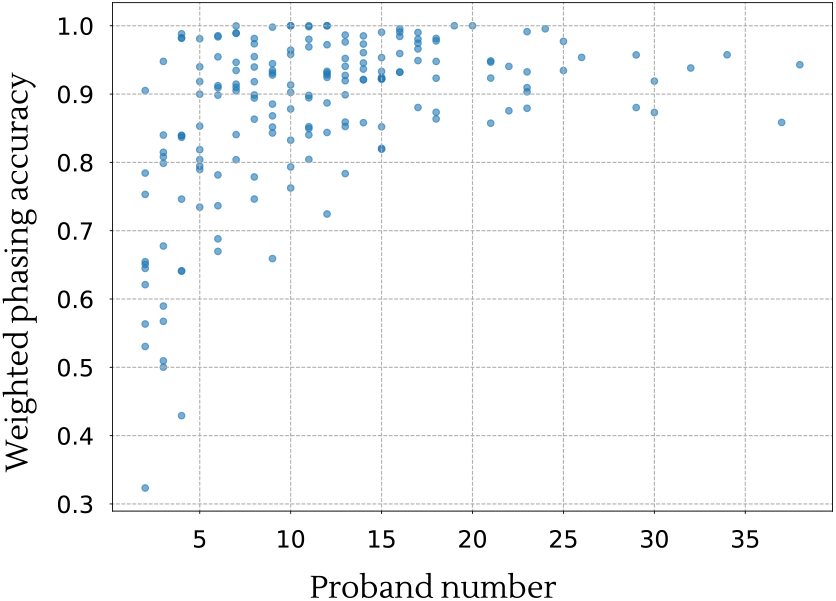
Phasing accuracy increases with the proband number in simulations. Each dot represents the weighted phasing accuracy of a clade alignment. The weighted phasing accuracy is calculated as the sum of proband phasing correctness ratio multiplied by the vertex alignment posterior probability.

### Dealing with errors

Realistic data is expected to feature errors in both ARG reconstruction and pedigree. While we would want eventually to modify the algorithm to handle both types of errors explicitly, the observation that the algorithm often provides multiple solutions near the oracle solution suggests a degree of robustness to perturbations – i.e, there are often some alternative ancestry paths available if the correct path is not. We will therefore first investigate the performance of the original algorithm on data with simulated errors, then discuss approaches to account for errors more formally.

#### Pedigree errors

Pedigree data may not reflect genetic transmissions due to adoptions, extra-pair paternity, or clerical errors. Based on y chromosome analysis, we found a rate of less than 1% mismatch on the paternal lineage, including all types of errors (Martínez 2025). Under the assumption that extra-pair paternity is a dominant source of mismatch, we randomly modified the BALSAC pedigree by introducing missing paternity errors.

Specifically, for each individual in BALSAC, we replaced the documented father with another father from the same generation with probability *λ* = 0.01. Here, generation number was estimated as the length of a longest path to a proband.

We then simulated 91 Balsac pedigrees with the missed parentage errors, and performed alignments on all simulated clade x pedigree combinations.

Figure 8 shows the results classified according to agreement between inferred pedigree assignments for the tree MRCA (i.e. root) of the clade relative to the simulated (oracle) assignment for the root, under the following 5 scenarios:

**Figure 8.**
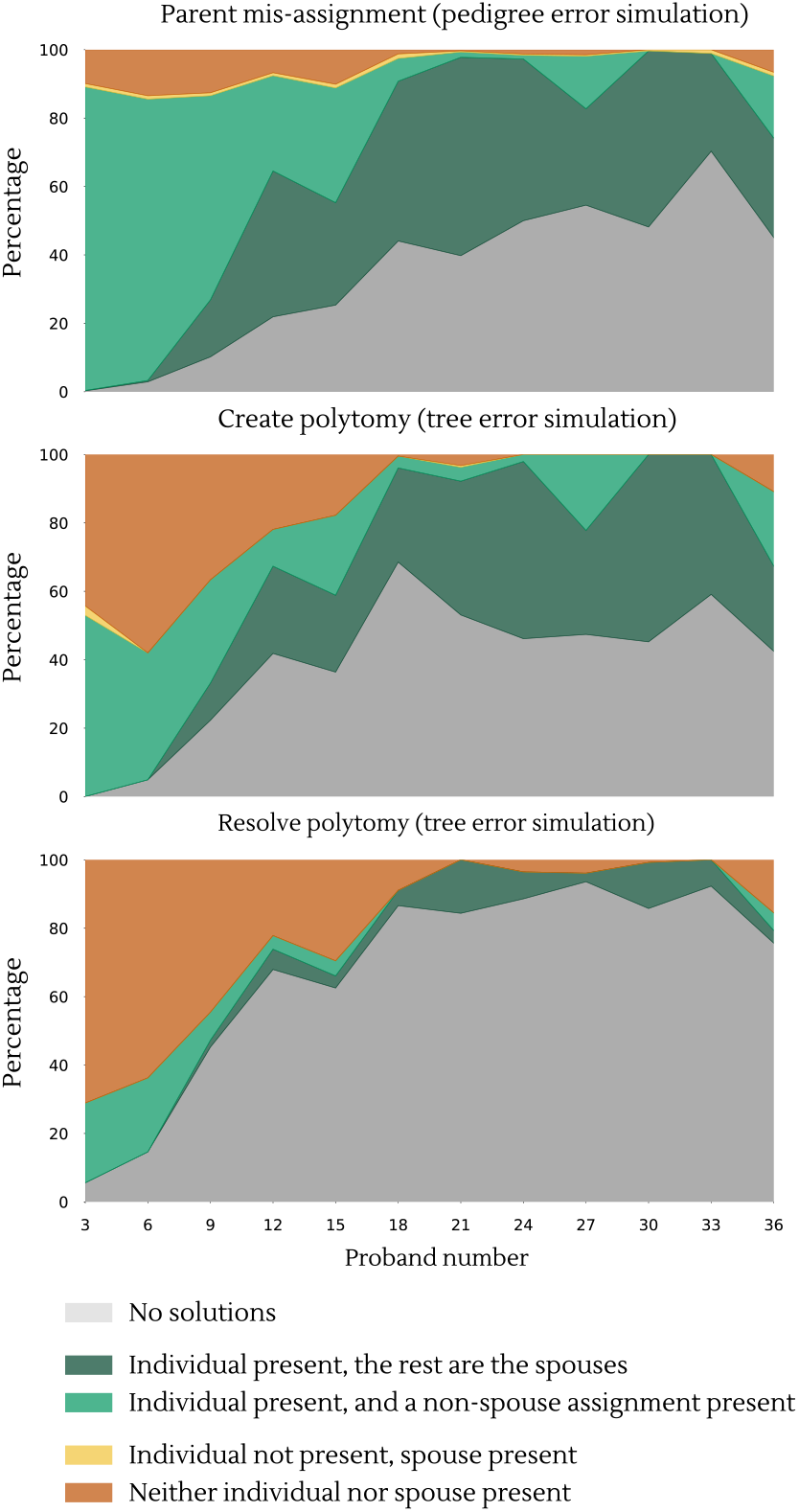
Correctness of tree MRCA inference under different error modes for the clades from Simulation setup. Results are shown within bins of size 3. Notice that there are only two clades with more than 36 probands (see Figure S3).

- No consistent assignments exist
- Correct individual found (i.e., the oracle ancestor is a feasible ancestor)
  (a) The remaining assignments are the oracle ancestor’s spouse(s).
  (b) There is also one or more assignment that isn’t the oracle ancestor’s spouse.
- Correct individual not found (i.e., the oracle ancestor is not a feasible ancestor):
  (a) One oracle ancestor’s spouse is a possible assignment.
  (b) All of the assignments are neither the oracle ancestor, nor any of their spouses.

Scenario 3(b) is the most problematic, as it may lead to incorrect conclusions about the origin of the allele.

#### Tree errors

We can use a similar approach for simulating tree errors. Unlike with pedigree errors, the types of errors we can expect to encounter have not been as extensively studied. Since our approach primarily focuses on the tree topology, we focus on the simplest type of topological error involving polytomies. Concretely, a tree polytomy might represent two close coalescent events and vice versa.

In order to test the algorithm’s stability in these cases, we performed similar simulations as in the previous section. Specifically, for every simulated clade, every possible modified clade which can be obtained by creating or resolving a polytomy was created.

Then, we aligned every obtained clade with the initial pedigree and classified the results by the possible clade root’s assignments (see Figure 8). The algorithm is more sensitive to tree errors than to pedigree errors. When the number of probands is large, there are often no valid alignments.

#### Maximum alignable subclade

One possible way to address scenario 1 (no alignment found) is to relax the constraints by solving the alignment problem for a subclade of the tree.

In this experiment, we used the same data with simulated tree errors. However, whenever the algorithm failed to find a solution, we considered all subclades of the tree that can be obtained by cutting one edge with at least 8 probands. We then selected the solutions for a subclade with the largest number of probands (see Figure 9). If there were multiple such clades, we chose one at random.

**Figure 9.**
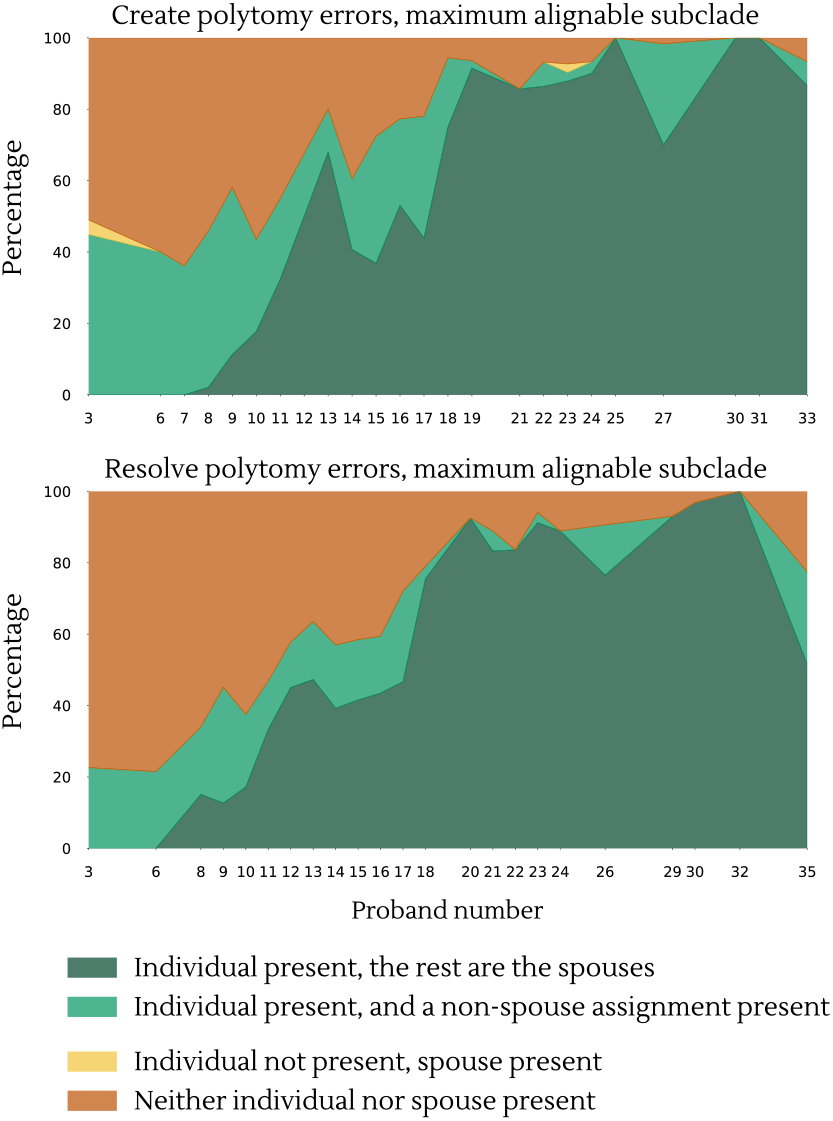
Correctness of tree MRCA inference using maximal clade alignment for the clades from Simulation setup, binned in groups of 30 alignable subclades.

From this section, we conclude that as long as the pedigree and inferred tree are sufficiently accurate, the algorithm is likely to provide informative solutions.

### Testing algorithm’s robustness on real data

Our original plan was to run the algorithm on ARGs inferred from CARTaGENE participants, since we have inferred ARGs for the entire cohort, and thus have ARG-defined clades for many rare, clinically-relevant variants (McClelland *et al*. (2025); Mejia-Garcia *et al*. (2025).) Unfortunately, we found that the inferred ARGs were often not topologically well-constrained – different runs of the ARG inference software ARG-needle would give comparable times for the TMRCA Mejia-Garcia *et al*. (2025), but quite different topologies. Because the founder event occurred many generations ago, many of the ancestral trees are star-like, with long terminal branches and short internal branches. Reconstructing to topology of such trees is difficult, since there is little information about the relative order of the splits. Performing comprehensive alignments for such star-like ARGs will require more detailed modelling of ARG uncertainty, which we will leave to future work since most ARG inference software does not provide a succinct way to encode the uncertainty.

To identify clades with a better-constrained local tree, we turned to a clinical study of Myotonic dystrophy type 1 (DM1). DM1 is an autosomal dominant inherited disorder and is the most common form of muscular dystrophy in adults (Harper 2009; Theadom et al. 2015). The age of symptom onset varies, from birth in congenital cases to 40+ for the late-onset form. Skeletal muscle weakness and myotonia are cardinal features of the disease, but additional symptoms exhibited by each patient, severity, and progression of the disease are variable. DM1 is caused by the expansion of a CTG trinucleotide repeat in the non-coding region of the DMPK gene. Despite large interindividual variations, the length of the CTG repeat is a rough determinant of DM1 severity (De Antonio et al. 2016).

The Saguenay-Lac-Saint-Jean (SLSJ) region of Quebec harbours many individuals of French-Canadian ancestry and has the highest incidence of DM1 worldwide, with a frequency of ≈ 1/6304. This region has undergone a strong founder effect, and consequently, most of the DM1 patients have one of two common haplogroups. Here, we focus on one of these two haplogroups. (Yotova et al. 2005). Since recruitment was phenotype-based, we expect to find a range of relatedness that may be more difficult to achieve in a population-based cohort like CARTaGENE.

### Cohort description

As part of a longitudinal study, the Groupe de recherche interdisciplinaire sur les maladies neuromusculaires (GRIMN) assembled a large, well-characterized familial cohort of more than 200 DM1 patients (Gagnon et al. 2018). Repeat length was measured, and abnormal expansion was confirmed for all patients and thoroughly analyzed (Overend et al. 2019). These patients also underwent genealogical reconstruction using the BALSAC database (Vézina and Bournival 2020) alongside whole-genome genotyping on the Infinium GSAv3.0 chip. Genotypes were cleaned using PLINK v1.9 (Purcell et al. 2007), ensuring individuals with at least 95% genotypes among all SNPs were retained. At the SNP level, we retained SNPs with at least 95% genotypes among all individuals, located on the autosomes and in Hardy–Weinberg equilibrium *p >m* 10^−6^. Cleaned genotypes were phased using Beagle v5.1 (Browning and Browning 2007). Phased genotypes were used to define 17 Mb haplotypes around the DM1 repeat expansion on chromosome 19 between positions 35571482 and 52984708 (see Figure 10) using the geneHapR R package (Zhang et al. 2023b). Haplotypes carrying the DM1 expansion were identified according to identity-by-descent (IBD) sharing measured using refinedIBD v17Jan20 (Browning and Browning 2013). We focused on families carrying the less common DM1 haplotype (Yotova et al. 2005). To validate parent-of-origin analyses, we selected as probands one individual from most families, defined by the clinic as groups of individuals sharing a parent or grand-parent. Other known cases were not used in the inference, but served to validate parent-of-origin analyses. We accidentally selected two individuals from one family (family 11) in this process, but decided to leave them in because the inclusion of close relatives does not affect the method.

**Figure 10.**
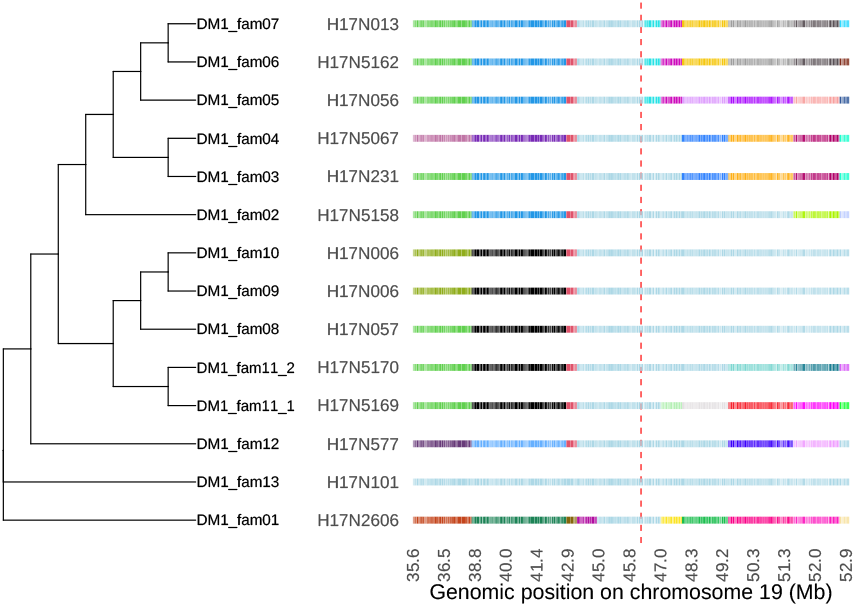
Phylogenetic tree based on haplotype reconstructions. The phylogenetic tree was built on 17 Mb haplotypes surrounding the DM1 expansion on chromosome 19. Haplotypes are shown on the y-axis and chromosomal positions on the x-axis. Instead of individual variant positions, linkage disequilibrium (LD) blocks are displayed. Haplotypes were segmented into variable-length segments, and a uniform color was applied across all haplotypes for segments where the LD blocks were consistently identical. The reference haplotype was randomly selected from one of the outgroup individuals.

Figure 10 shows the haplotype structure for these individuals around the DM1 locus, and shows long stretches of shared haplotypes among subsets of carriers. Given the clear haplotype structure, we reconstructed the local tree using RAxML-NG v1.2.0 (Stamatakis 2014) with the GTR+G model and default parameters to build phylogenetic trees on unique haplotypes in families. The best tree was selected according to visual inspection of LD blocks inferred using haploview v4.2 (Barrett *et al*. 2005).

### Alignment results

The best gene tree was aligned to the genealogical tree for the same probands. In total, 32 possible vertex alignments were inferred by the algorithm, with posterior probabilities ranging from 1.68% to 5.01%. One of the obtained edge alignments is depicted in Figure 11. It shows the potential pedigree paths through which the mutation was inherited. Additionally, the vertex inclusion probabilities (averaged over all alignments) are shown for each displayed vertex. Carrier statuses of ancestors were not used in the alignments, providing a way to evaluate the accuracy of the parent-of-origin estimates – we find that the parents-of-origin estimated with high posterior probability all match the clinical record, whereas parents-of-origin with uninformative posteriors are, as expected, poorly determined. A single couple was found as a compatible common ancestor to the set of probands. This couple was married in 1638 in the Perche region of France, and is one of the largest contributing couples to the SLSJ region. The couple is thus a plausible candidate for contributing a founder variant found at appreciable frequency in the region, but the high contribution also makes them more likely to be false positive if there is a problem with the inferred tree. Two branches in the gene tree (DM1 fam01 and DM1 fam13) share only short haplotypes with other nodes, and their precise position in the tree is thus poorly constrained. Excluding the two corresponding branches leaves five additional potential ancestral groups that could have introduced the variant into the population, so that we cannot be certain about the identity of the MRCA.

**Figure 11.**
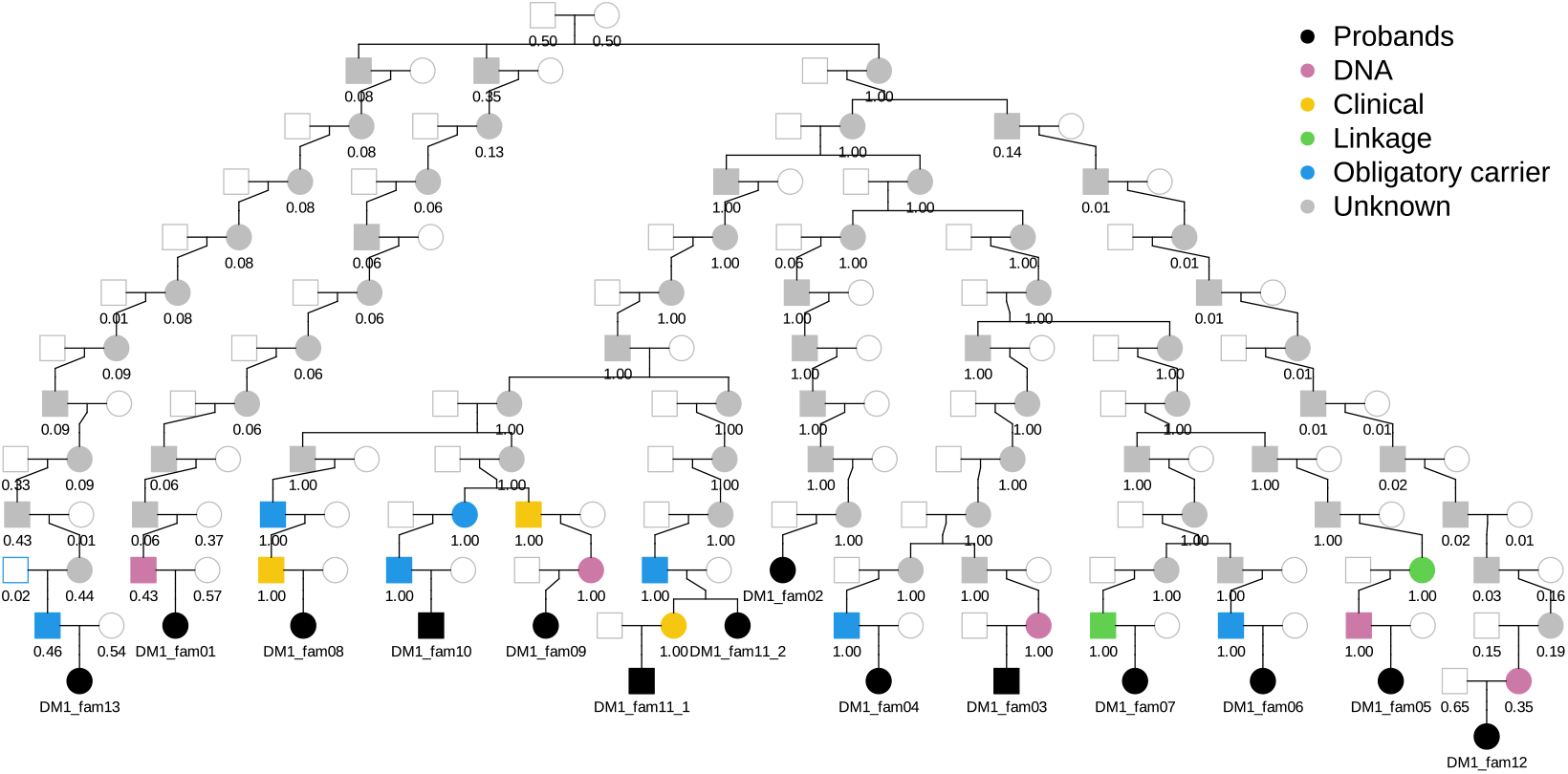
Genealogical path from the putative founder carrying the DM1 variant to affected descendants. Filled circles or squares indicate carrier ancestors, colored according to the clinical method of diagnosis when available. Ancestors shown in blue were not genetically or clinically tested; however, they are considered obligatory carriers due to having affected offspring and siblings. Alignment was performed in the unphased regime for the 13 leaves of the tree only.

### Further work

#### Dealing with errors

The approach that we’ve developed works correctly with perfect data. However, as we can see, the algorithm can be sensitive to various types of errors in the input data. In particular, we observed that even small changes in the tree can result in the algorithm finding only wrong solutions or finding no solutions at all.

Future work could focus on systematically characterizing the possible types of errors — especially those affecting the tree structure — and on extending the proposed approach to improve its robustness against such errors.

#### Solving the entire ARG

So far, our efforts have focused solely on aligning a single local tree with the pedigree. The natural next step would be to align the pedigree with the entire ARG. That is, instead of finding all the solutions consistent only with one tree, the resulting set of solutions would need to satisfy the topologies of all trees simultaneously.

Using this information, we can significantly reduce the set of resulting solutions, since we have more constraints to consider.

On the other hand, the increased amount of information may also amplify the impact of errors, which signifies the importance of detecting and addressing such errors in real data.

#### Occasional long inference time for edge alignments

Although the alignment problem appears to be relatively well-constrained for most inputs we have worked with, there are cases where the input data is poorly constrained, leading to significantly longer search times. Specifically, this happened when the algorithm was used to search for all possible edge alignments, which turned out to be extremely slow due to a high variability. On the other hand, the running time on the same input was relatively low when only one edge alignment was found for each vertex alignment.

Nevertheless, one possible improvement would be to optimize the running time of the developed algorithm in these scenarios.

## Data availability

The code for the developed algorithm, as well as the code for simulating errors can be found [here](https://github.com/gravelLab/pedigree-ARG-matching). The BALSAC and CARTaGENE datasets can be obtained via the corresponding procedures.

## Acknowledgments

We would like to thank Sergiy Kozerenko for his insightful comments, which helped improve the earlier versions of this paper. We also thank Shadi Zabad, Marc Henein, Alejandro Mejia-Garcia, Marianne de Vriendt, Ivan Krukov, and Andjela Todorovic for useful discussions and early explorations of this problem.

## Appendix A

### Validating an alignment

By using the climbing strategy described in Step (b) of the alignment algorithm, we find a set of candidate alignments between the tree and the pedigree. To save on computational time, we did not ensure that the paths from probands to the common ancestor were fully disjoint. This condition is validated a posteriori.

In this section, we are going to formally define what a valid alignment is and develop an algorithm for performing the validation effectively.

#### Preliminaries

We first introduce some general notation. Here and further, all the considered graphs are simple (no loops [i.e., edges from a node to itself] or multiple edges between a pair of nodes are allowed). Suppose we have a directed graph *D* = (*V*(*D*), *E*(*D*)) with *V*(*D*) being the set of *vertices* and *E*(*D*) being the set of *directed edges* (or *arcs*). We can use this model to represent either pedigree or genetic ancestry relationships, with directed edges *uv* pointing from a child vertex *u* to a parent vertex *v*. In a pedigree, the directed edge represents a parent-offspring relationship, whereas in a gene tree, these represent ancestor-descendant relationships.

For a vertex *u* ∈ *V*(*D*), the set *N*^−^(*u*) is called the *in-neighborhood* of *u*: *N*^−^(*u*) = {*v* | *vu* ∈ *E*(*D*)}. Similarly, *N*^+^(*u*) = {*v* | *uv* ∈ *E*(*D*)} is the *out-neighborhood* of *u*. A (directed) *walk* is a sequence of vertices *u*_1_, …, *u*_*n*_ such that for all 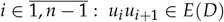 is an arc. A (directed) *path* is a walk in which all the vertices are distinct. Given two vertices *u, v* ∈ *V*(*D*), denote by 𝒫_*uv*_ the set of all paths from *u* to *v*. That is,

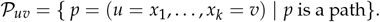

Similarly, we can define the set of all paths in the following way:

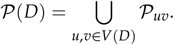

A *tree* is a connected acyclic graph. A *directed tree* is a DAG whose undirected version is a tree. A *directed rooted tree T* is a directed tree where one vertex *r*(*T*) has been selected as the root. An *in-tree* is a directed rooted tree where all the arcs point towards the root. Finally, a tree *T* is said to be *non-unary* if every non-leaf vertex has at least two children. Coalescent trees can be represented as non-unary in-trees.

Notice that every non-root vertex *v* in an in-tree has a unique parent, referred to as *π*_*T*_ (*v*). Thus *π*_*T*_ (*v*) is the vertex connected to *v* on the path to the root *r*(*T*). A *sibling* of a vertex *v* is another vertex in *T* who shares a parent with *v*. A *descendant vertex* of *v* is a vertex in *V*(*D*) that is either a *v*’s child or is a descendant of one of *v*’s children. In other words, a vertex *u* descends from *v* if 𝒫_*uv*_ ≠ Ø. For a vertex *v* ∈ *V*(*D*), let *d*_*D*_ (*v*) denote the list of all the *v*’s descendants. A *leaf vertex* is a vertex with no descendants. Non-leaf vertices are called *internal vertices*.

In the context of our problem, a pedigree is simply a DAG. As before, we denote the pedigree by *P*, the coalescent tree by *T* where *L*_*T*_ is the set of leaf vertices in *T*. Now, we prove a lemma that we will need later.

##### Lemma 1. For a directed tree T with n vertices, we have

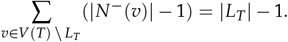

*Proof*. By the handshaking lemma for directed graphs, we have

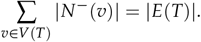

Since *T* is a tree, |*E*(*T*)| = |*V*(*T*)| − 1. Hence,

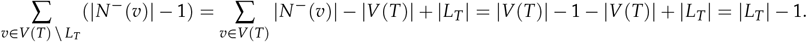

##### Corollary 1.

*For a vertex u in a directed tree T, the following holds:*

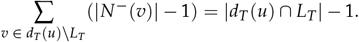

*Proof*. Consider the subtree *T*_*u*_ descending from *u* with *u* being the root and apply Lemma 1.

#### From diploid to haploid pedigree

The input pedigree specifies relations between diploid individuals. However, the algorithm has to assign gene tree vertices to either the maternally-inherited or the paternally-inherited ploids in a pedigree individual. We therefore build a *ploid pedigree* before starting the alignment procedure.

To build the ploid pedigree, we create two vertices for every individual in the pedigree, each representing a ploid. An individual’s paternally-inherited ploid is connected to the father’s ploids and the maternally inherited ploid to the mother’s ploids. The resulting ploid pedigree captures all possible paths through which genetic material may have been transmitted under a standard diploid transmission model.

#### Formal problem definition

As mentioned in Problem statement, our goal is to find the set of all the valid histories of genetic transmissions that respect *P* and *T*. This concept can be formalized by defining the notion of the *edge alignment*, in which each edge from the gene tree is assigned a path in the pedigree, such that intersections occur only at common ancestors. Formally:

##### Definition 1.

*An* ***edge alignment*** *between a pedigree P and a coalescent tree T is a function* Γ : *E*(*T*) → 𝒫 (*P*) *such that:*

- ∀*u* ∈ *V*(*T*), ∀*v, w* ∈ *N*^−^(*u*) : Γ(*vu*) = (*x*_1_, …, *x*_*m*_), Γ(*wu*) = (*y*_1_, …, *y*_*k*_) =⇒ *x*_*m*_ = *y*_*k*_, Γ(*vu*) ∩ Γ(*wu*) = {*x*_*m*_ };
- ∀*u* ∈ *V*(*T*), *u* ≠ *r*(*T*), ∀*v* ∈ *N*^−^(*u*) : Γ(*vu*) = (*x*_1_, …, *x*_*m*_), Γ(*uπ*_*T*_ (*u*)) = (*y*_1_, …, *y*_*k*_) =⇒ *x*_*m*_ = *y*_1_, Γ(*vu*) ∩ Γ(*uπ*_*T*_ (*u*)) = {*x*_*m*_ }.

Since vertices in gene trees correspond to most recent common ancestors (MRCAs), pedigree paths can meet at most at one position corresponding to that MRCA. We say that a pedigree vertex *u* ∈ *V*(*P*) *belongs* to a map Γ : *E*(*T*) → 𝒫 (*P*) if there exists an arc *e* ∈ *E*(*T*) such that *u* ∈ Γ(*e*).

An edge alignment captures the full transmission history, but we can also develop a simplified concept to capture the MRCAs induced by an edge alignment – a *vertex alignment*. A vertex alignment is simply a function *f* : *V*(*T*) → *V*(*P*) that maps gene tree nodes to pedigree nodes. For a vertex *x* ∈ *V*(*P*), we say that *x* is a *coalescent vertex candidate* for a vertex alignment *f* if there exists *u* ∈ *V*(*T*) where *f* (*u*) = *x*.

An edge alignment implies a vertex alignment. Indeed, if we want to find the pedigree vertex for *t* ∈ *V*(*T*), we can simply put *f* (*t*) = *x*_1_ where Γ(*tπ*_*T*_ (*t*)) = (*x*_1_, …, *x*_*k*_). This relation is formalized in the next definition.

##### Definition 2.

*For a vertex alignment f between a coalescent tree T and a pedigree P, an edge alignment* Γ *is said to* ***correspond*** *to f if*

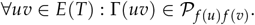

##### Corollary 2.

*If an edge alignment* Γ *corresponds to a vertex alignment f, then the conditions stated in Definition 1 can be reformulated as:*

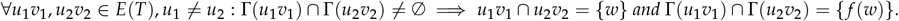

##### Corollary 3.

*Given a vertex alignment f* : *V*(*T*) → *V*(*P*) *and a corresponding edge alignment* Γ, *for a pedigree vertex that belongs to* Γ, *but is not a coalescent vertex candidate for f, there exists a unique path to which it belongs*.

Finally, for a vertex alignment *f*, we will use F _*f*_ to denote the set of all the valid edge alignments that correspond to *f*. This allows us to define what a *valid* vertex alignment is.

##### Definition 3.

*A vertex alignment f is* ***valid*** *if* ℱ _*f*_ ≠ Ø.

#### Single MRCA case

Before we develop an approach that can work with the whole alignment, we’re going to solve a sub-problem involving only one MRCA. That is, given a tree vertex *u* with *N*^−^(*u*) = {*u*_1_, …, *u*_*n*_ }, we need to find out whether there are paths from 𝒫 _*f* (*u*1) *f* (*u*)_, …, 𝒫 _*f* (*un*) *f* (*u*)_ that intersect in *f* (*u*):

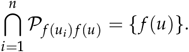

This a common algorithmic problem that is usually modelled as a maximum flow problem, and can be solved using the Ford–Fulkerson algorithm (Ford and Fulkerson 1956).

#### Maximum flow problem

A *network* (*G, c*) is a directed graph *G* with a capacity function *c* : *E*(*G*) → ℝ^+^ defined on its arcs. A *flow network* (*G, s, t, c*) is a network with two distinct distinguished vertices *s, t* ∈ *V*(*G*) called the *source* and the *sink*, respectively (Williamson 2019).

Flow networks can be used to represent, e.g., water going through a pipe network. We found that they were also useful to represent the flow of ancestry through a pedigree.

A flow is a map *h* : *E*(*G*) → ℝ^+^ that satisfies the following constraints:

- **Capacity constraint** The flow must not exceed the capacity: ∀*uv* ∈ *E*(*G*) : *h*(*uv*) ≤ *c*(*uv*);
- **Conservation of flows** Except for the source and the sink, the sum of the flows entering a vertex must be the same as the sum of the flows leaving the vertex:

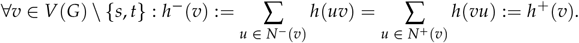

Here, for a vertex *v*, the values *h*^−^(*v*) and *h*^+^(*v*) are called the *in-flow* and the *out-flow* of a vertex, respectively. The value |*h*| of the flow *h* is the flow reaching target vertex *t*:

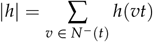

Finally, the maximum flow is the flow *h* with maximum value |*h*|. Multiple algorithms can solve this problem, such as the Ford–Fulkerson algorithm, the Edmonds-Karp algorithm, etc. (Williamson 2019).

#### Validating that paths are edge-disjoint

We can already use the maximum flow problem to verify a necessary condition for an alignment to be valid. Specifically, we can check whether all the corresponding pedigree paths are edge-disjoint. We can do that by building a supplementary graph *G*_*f* (*u*)_ from *P* to verify that condition. Here, *u* is a non-leaf vertex in *T* with *N*^−^(*u*) = {*u*_1_, …, *u*_*n*_ }.

Start with *P*_*f* (*u*)_ — the ascending pedigree from *f* (*u*_1_), …, *f* (*u*_*n*_) to *f* (*u*) (that is, the minimal subgraph of *P* containing all paths from *f* (*u*_1_), …, *f* (*u*_*n*_) to *f* (*u*)). Then, add the source and target vertices *s, t* to it. Now, for all 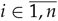 add arcs *s f* (*u*_*i*_). Next, add the edge *f* (*u*)*t*. Denote the resulting graph by *G*_*f* (*u*)_ (see Figure 12).

**Figure 12.**
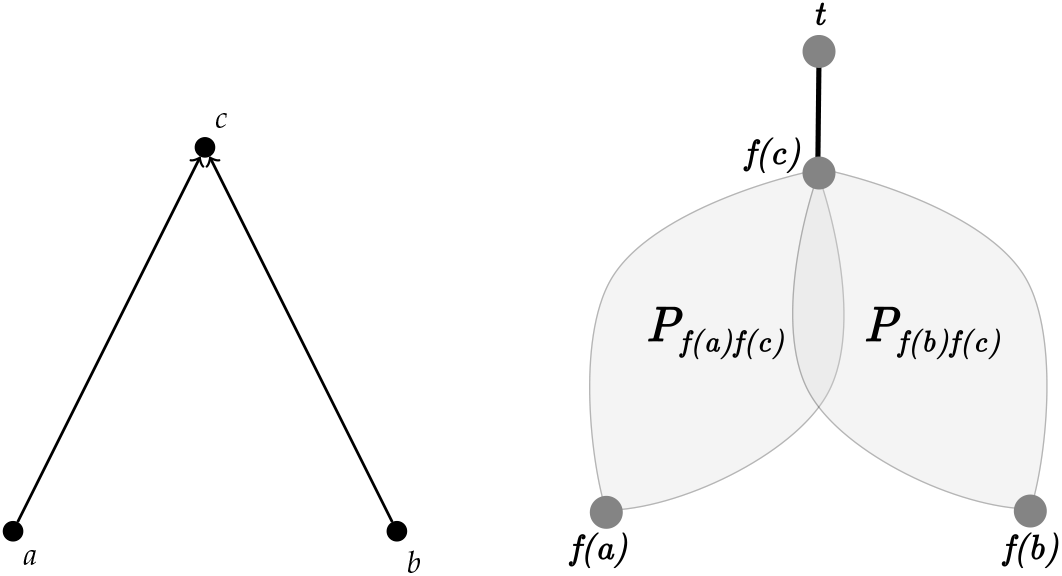
An illustration of the constructed graph *G*_*f* (*c*)_ that can be used to identify a series of edge-disjoint paths for a coalescent event with two child nodes.

Finally, assign every directed edge the capacity of 1: ∀*e* ∈ *E*(*G*_*f* (*u*)_) : *c*_1_(*e*) = 1 where *e* ?= *f* (*u*)*t*. For *f* (*u*)*t*, we have *c*_1_(*f* (*u*)*t*) = *n*.

To check whether the paths are edge-disjoint, we can simply run a maximum flow algorithm on the built flow network *N*_1_ = (*G*_*f* (*u*)_, *s, t, c*_1_) and verify whether the network flow is equal to *n*. If so, then the paths are edge-disjoint. Otherwise, the alignment is not valid (Cormen et al. 2022).

#### Validating that paths are vertex-disjoint

Edge-disjoint paths could still share a vertex. To ensure that the paths are also vertex-disjoint, we need a slightly more complex graph. This can be done by building a new flow network *N*_2_ from *N*_1_.

Start with *G*_*f* (*u*)_. Next, for every vertex *v* ∈ *V*(*G*_*f* (*u*)_) \ {*s, t*}, add a new vertex *v*^′^ with an edge *v*^′^ *v* and a capacity of 1. Here, *v*^′^ is called the **access vertex** of *v*. Then, we want to reconnect all the vertices that were previously connected with *v* to *v*^′^. Formally speaking, we will remove every arc *uv* ∈ *E*(*G*_*f* (*u*)_) and add a new arc *uv*^′^ instead with a capacity of 1. An example of this operation is shown in Figure 13. In other words, all the directed edges that were previously going to a vertex *u* directly now will have to pass through its access vertex *u*^′^ before reaching it. Finally, add an arc *f* (*r*(*T*)^′^ *t*) with a capacity of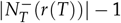.

**Figure 13.**
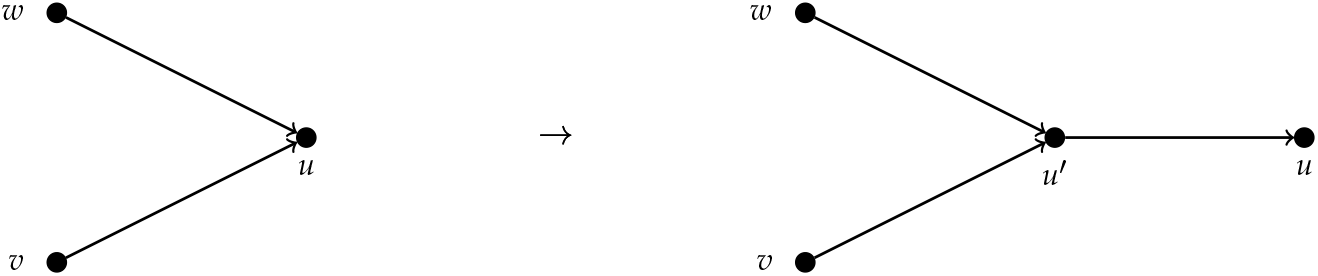
Illustration of the operation of adding an access vertex to ensure that we identify edge-disjoint paths.

Denote the resulting graph as *K*_*f* (*u*)_ and put *N*_2_ = (*K*_*f* (*u*)_, *s, t, c*_2_). Now, the problem of finding a series of vertex-disjoint paths in *G*_*f* (*u*)_ is equivalent to finding a series of edge-disjoint paths in *K*_*f* (*u*)_. We can use the developed approach to verify whether *u* is a valid MRCA for *u*_1_, …, *u*_*n*_.

#### Solving the general case

We can now build on the approach developed for the single MRCA case to verify the correctness of the whole alignment. To begin with, we use the same procedure to build the graph *H*_*f*_ by taking the union of graphs *K*_*f* (*u*)_ for every non-leaf coalescent vertex *u* in *T*. Specifically,

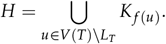

An example of the graph *H* is shown in Figure 14.

**Figure 14.**
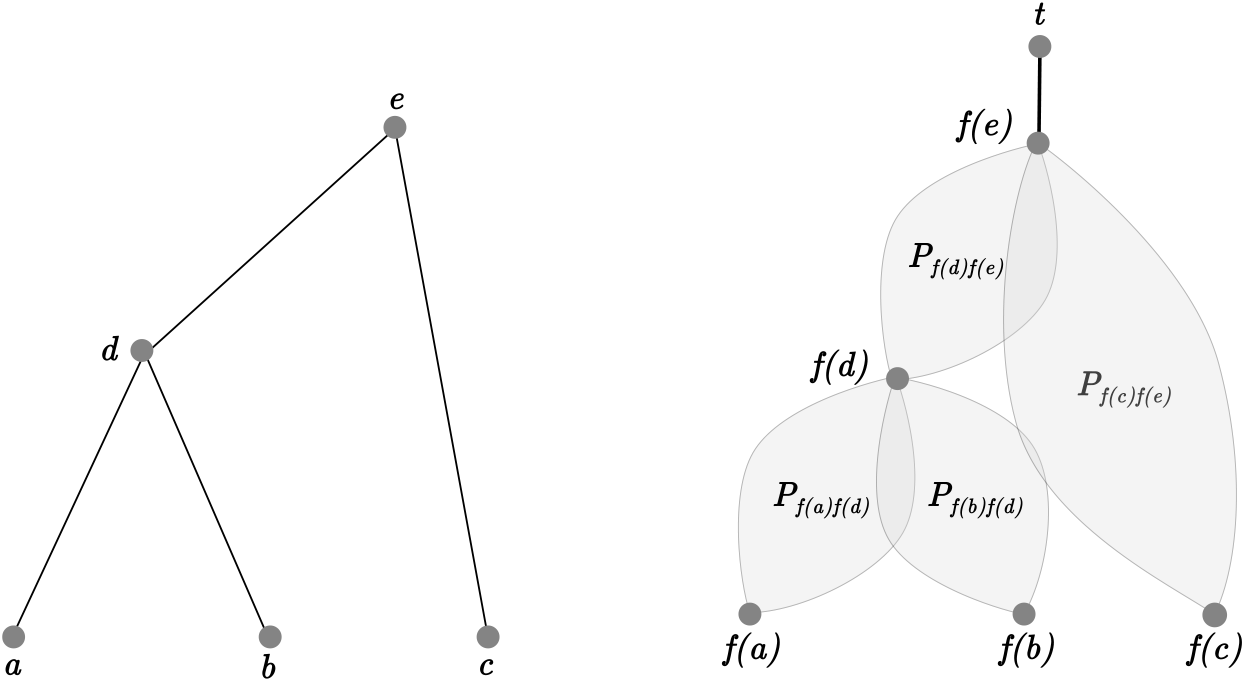
An example of a coalescent tree and the corresponding graph *H*_*f*_, constructed using the tree, the pedigree, and a vertex alignment *f*.

For a valid vertex alignment *f* and a corresponding edge alignment Γ ∈ ℱ _*f*_, we can always construct an auxiliary path map *k*_*p*_ : *E*(*T*) → 𝒫 (*H*_*f*_) that follows the additional arcs and access vertices. That is, for an arc *e* ∈ *E*(*T*) and its corresponding pedigree path Γ(*e*) = (*x*_1_, …, *x*_*k*_), we have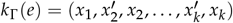.

Finally, let *α* : *V*(*H*_*f*_) \ {*s, t*} → *V*(*P*) be the function that returns the non-access version of a vertex in *H*_*f*_. That is, if *x* ∈ *V*(*H*_*f*_) isn’t an access vertex, then *α*(*x*) = *x*. Otherwise, if *x* is an access vertex of some other vertex *y*, then *α*(*x*) = *y*.

Now, we can solve the maximum flow problem for *H* to obtain a necessary condition for a valid vertex alignment.

##### Theorem 1.

*If a vertex alignment f between T and P is valid, then the maximum flow of the graph H* = *H*_*f*_ *passing from s to t is equal to* |*L*_*T*_ |.

*Proof*. Given the outgoing arcs’ capacities from the source, the flow cannot be larger than |*L*_*T*_ |. We will therefore proceed by showing the existence of a flow with flow value |*L*_*T*_ |, which will therefore be a maximizing flow.

For brevity, let *r* = *r*(*T*) denote the root of the coalescent tree. Since *f* is a valid vertex alignment, ℱ _*f*_ ≠ Ø. Let Γ ∈ ℱ _*f*_ be a corresponding edge alignment. Then, let *k* = *k*_Γ_ be the auxiliary path map. Let’s define the flow *h* the following way:

- The flow coming from the source to each of the leaf vertices’ pedigree candidates is equal to 1:

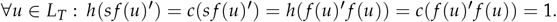
- For every arc in the gene tree, the corresponding path’s arcs have the flow equal to their capacity:

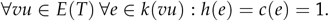
- The flow of an arc going from a coalescent vertex candidate’s access vertex to the target vertex is equal to its capacity:

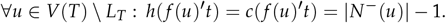
- The flow going from the root’s candidate to the target is 1:

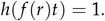
- For the rest of the arcs, the flow is 0.

Let *x* be a vertex belonging to *k* with *α*(*x*) ≠ *f* (*u*) for all vertices *u* in *T*. That is, *x* is neither a coalescent vertex candidate nor an access vertex of such a vertex. Based on the construction of *h*, notice that for every such vertex *x*, there exist unique in-going and out-going edges with a non-zero flow.

Since *x* belongs to *k*, by Corollary 3, there exists *e* ∈ *E*(*T*) with *x* ∈ *k*(*e*). Moreover, *k*(*e*) = (*x*_1_, …, *y, x, z*, …, *x*_*m*_^′^, *x*_*m*_) and *h*(*yx*) = *h*(*xz*) = 1. Now, we want to show that *y* and *z* are unique. Suppose there exists a vertex *z*_1_ ≠ *z* with *h*(*xz*_1_) = 1. This implies that there exists another edge *e*_1_ ∈ *E*(*T*) with *x* ∈ *k*(*e*_1_), which contradicts that Γ is an edge alignment as *x* ∈ *k*(*e*) ∩ *k*(*e*_1_) and *x ≠ x*_1_, *x*_*m*_^′^, *x*_*m*_. We can prove that *y* is unique in the same way. For such a vertex *x*, put *ν*(*x*) = *z* and *µ*(*x*) = *y*.

Now we can show that *h* is a valid flow with |*h*| = *m*.

##### Capacity constraint

Notice that if the flow of an arc is non-zero, then it’s equal to its capacity. Therefore, the capacity constraint is satisfied.

##### Conservation of flows

For every vertex *v* ∈ *V*(*H*) \ {*s, t*}, we need to verify that *h*^−^(*v*) = *h*^+^(*v*). Suppose, on the contrary, that there is such a vertex *x* for which it doesn’t hold.

**Case 0:** Vertices with no flow.

Notice that, by construction, for a vertex *v* ∈ *V*(*H*) with *v* ≠ *s, t*, if *α*(*v*) does not belong to *k*, then *h*^+^(*v*) = *h*^−^(*v*) = 0.

**Case 1:** *x* is an access vertex of a coalescent vertex candidate.

Let’s consider the case when *x* is an access vertex of a coalescent vertex candidate, that is *x* = *f* (*u*)^′^, *u* ∈ *V*(*T*). If *u* ∈ *L*_*T*_, then we have *h*^+^(*x*) = *h*(*x f* (*u*)) = 1 and *h*^−^(*x*) = *h*(*sx*) = 1.

Otherwise, we have *N*^+^(*x*) = {*t, α*(*x*)} and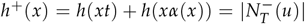. Since *k* is an edge alignment between *T* and *H*, for every child vertex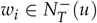, there is a pedigree path *k*(*w*_*i*_*u*) joining *f* (*w*_*i*_) and *f* (*u*) in *H*. Let *v*_*i*_ be the vertex on that path that is connected to *f* (*u*)^′^ in the pedigree. That is, *v*_*i*_ ∈ *k*(*w*_*i*_*u*) and *v*_*i*_ *f* (*u*)^′^ ∈ *E*(*H*). Notice that all the *v*_*i*_ are distinct, as *k* is an edge alignment, and *h*^+^(*v*_*i*_ *f* (*u*)^′^) = 1. Therefore,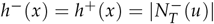.

**Case 2:** *x* is a coalescent vertex candidate.

That is, *x* = *f* (*u*), *u* ∈ *V*(*T*). Notice that, in this case, *h*^−^(*v*) = *h*(*v*^′^ *v*) = 1. If *v* = *f* (*r*), then *h*^+^(*v*) = *h*(*vt*) = 1. Otherwise, there always exists a unique vertex *w* with *vw* being an arc and *h*(*vw*) *>m* 0. Specifically, we have *k*(*uπ*_*T*_ (*u*)) = (*v, w*, …). By construction of *h*, we have *h*^+^(*v*) = *h*(*vw*) = 1.

**Case 3:** *α*(*x*) is not a coalescent vertex candidate.

Since *α*(*x*) is not a coalescent vertex candidate and it belongs to *k*, we can conclude that *x* also belongs to *k*. Moreover, we have *h*^+^(*x*) = *h*(*xµ*(*x*)) = 1, *h*^−^(*x*) = *h*(*ν*(*x*)*x*) = 1.

Therefore, *h* is a valid flow.

##### Flow value

Now we need to show that its flow value is *m* := |*L*_*T*_ |. By construction, *N*^−^(*t*) = { *f* (*u*)^′^ | *u* ∈ *V*(*T*) \ *L*_*T*_ } ∪ { *f* (*r*)}. Now, using Corollary 1, we have:

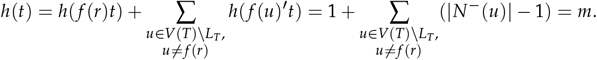

Finally, notice that this flow is maximum as

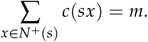

Therefore, *h* is a maximum flow with |*h*| = *m*.

Unfortunately, the converse is not true. That is, if the value of the maximum flow is equal to the number of leaves (probands) in the tree, this doesn’t imply that the alignment is valid. Specifically, the maximum flow doesn’t guarantee that the tree vertices coalesce at their MRCA (see Figure 15). A solution to this problem would be to use Multi-commodity Flow (Ahuja et al. 1993), where, for an arc *t*_1_*t*_2_ ∈ *E*(*T*), we form a commodity of demand 1 with *f* (*t*_1_), *f* (*t*_2_) being the sink and the source, respectively.

**Figure 15.**
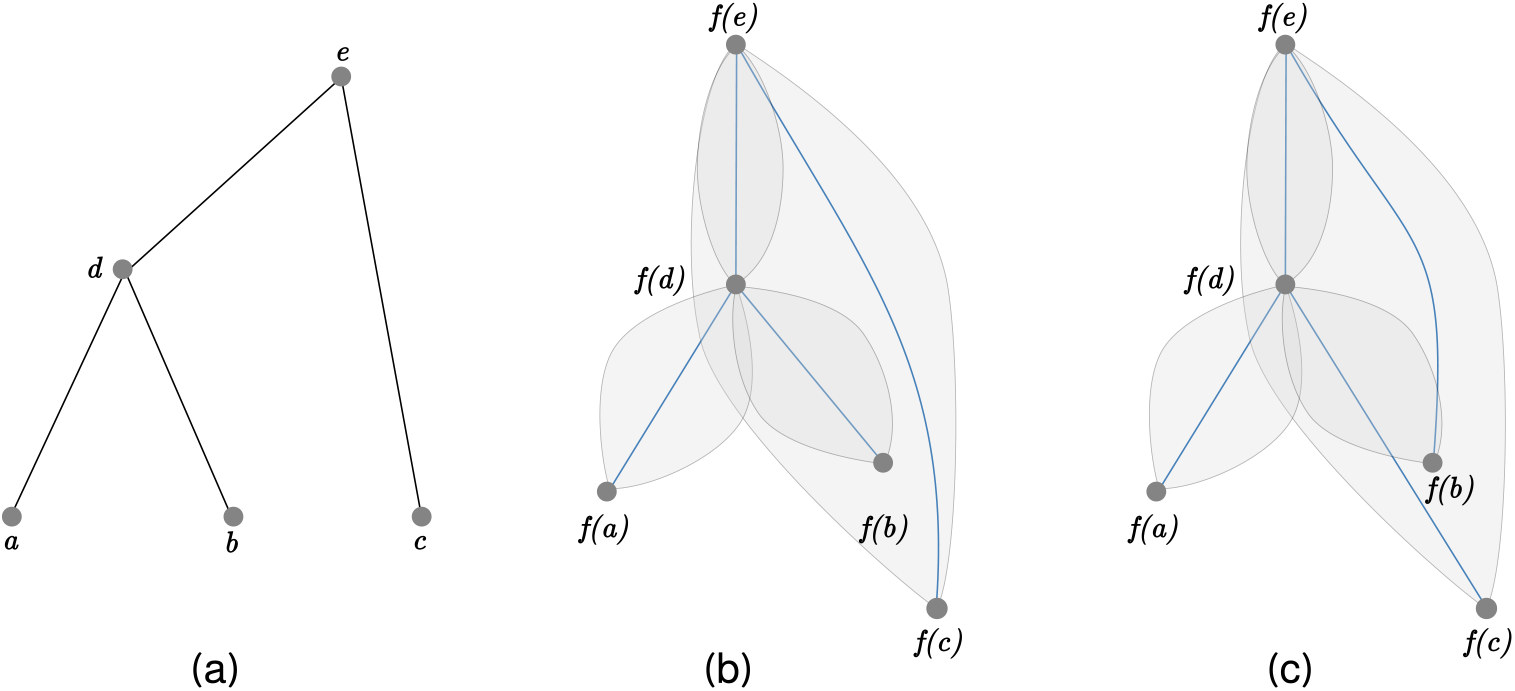
An example illustrating that maximum flow doesn’t always produce a valid alignment, where (a) represents the coalescent tree; (b) and (c) show maximum flows that produce valid and invalid edge alignments, respectively.

Alternatively, we can perform a simple verification to check whether the maximum flow yields a valid solution. We find that the alignment produced by the maximum flow is quite often correct. In the cases where it’s not true, we resort to a brute-force search to identify one (or all) possible edge alignments. This problem can be solved in a manner similar to the climbing strategy discussed in Appendix B (for instance, a simple brute-force search can be used).

## Appendix B

Andrii Serdiuk, et al. 19

### Climbing strategy

In this section, we’re going to describe the algorithm described in Algorithm description in more detail.

#### Step (a): Preprocessing

We want to traverse *T* in such a way that children vertices are traversed before their parents. To do this, we find a topological ordering of *T* (which exists since *T* is a DAG). We do this by ordering the vertices by their distance to the root (or to a proband).

#### Step (b): Climbing step

This is the main recursive step of the algorithm. Assuming that children of an inner vertex *u* ∈ *V*(*T*) already have candidate pedigree assignments, we want to find pedigree assignments for *u*. Because the algorithm is initiated with candidate pedigree assignments for the leaves of *T* (the proband), successive application of this recursion will provide assignments to all vertices in *T*.

The idea is that vertex *u* in the gene tree corresponds to a MRCA for its children vertices.

##### Definition 4.

*Let A* = {*a*_1_, …, *a*_*n*_ } *be a list of distinct pedigree vertices in P. A vertex a* ∈ *V*(*P*) *is called a* ***pedigree MRCA*** *for the vertices a*_1_, …, *a*_*n*_ *if there exists a map p*_*a*_ : *A* → 2^*V*(*P*)^ *such that:*

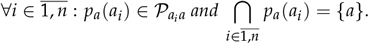

Thus the corresponding vertex *f* (*u*) in the pedigree must be a *pedigree MRCA* for the pedigree vertices assigned to children of *u*.

The pseudocode for the climbing step is shown in Algorithm 1.

##### Algorithm 1 Algorithm’s pseudocode

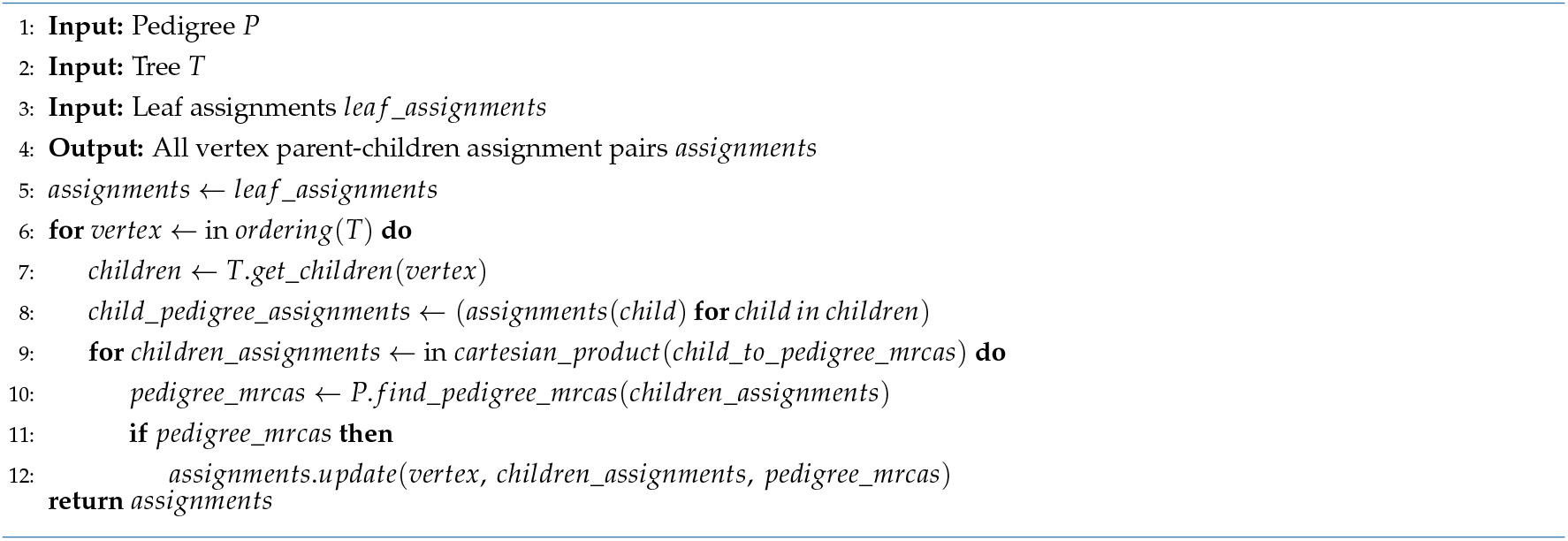

Our implementation differs from this at a few junctions. For example, we found that it was slightly faster to define a function *f ind*_*pedigree*_*MRCA*_*candidates*() that identifies a superset of the pedigree MRCAs, are rely on the maximum flow algorithm at step (c) to filter out incorrect MRCAs. Therefore, choosing the implementation for the *f ind*_*pedigree*_*mrca*_*candidates* function is a question of trade-off between faster local verification and a larger search space. This is a common issue arising in CSP problems (Russell and Norvig 2016).

Additionally, taking the Cartesian product of all the children assignments is highly inefficient for solving polytomies. We can use more efficient strategies that ensure that all possibilities have been accounted for. One example solution of this step is described in Climbing step implementations, below.

#### Step (c): Reconstruction

Having found the possible pedigree candidates for the tree vertices, we can recursively build the set of all the corresponding alignments between *T* and *P*. As pointed out before, the number of resulting alignments grows rapidly as the number of pedigree candidates for a vertex increases.

Next, we can use the steps developed in Appendix A to filter out invalid alignments from this set by trying to find at least one corresponding edge alignment. The resulting set of vertex alignments is the solution to our problem.

#### Climbing step implementations

In this section, we are going to give an example implementation of step (b). Firstly, let’s address the *f ind*_*pedigree*_*mrca*_*candidates* function that returns a superset of all the pedigree MRCAs for a set of pedigree vertices. We have two partially contradicting goals in mind:

- The running time of *f ind*_*pedigree*_*mrcas* should be low.
- The resulting solution set found should contain as few invalid assignments as possible.

In the current implementation, we developed a notion of a *potential pedigree MRCA* for a set of pedigree vertices.

##### Definition 5.

*Let A* = {*a*_1_, …, *a*_*n*_ } *be a list of distinct pedigree vertices in P. A vertex a* ∈ *V*(*P*) *is called a* ***potential pedigree MRCA*** *for the vertices a*_1_, …, *a*_*n*_ *if there exists an injection p* : *A* → *N*^−^(*a*) *such that there is a path from a to* Γ(*a*).

In other words, a vertex *a* is a potential pedigree MRCA for *A* if every vertex from *A* can reach *a* via a distinct child. Notice that the set of all the potential pedigree MRCAs is a superset of all the MRCAs. The intuition for this approach comes from the fact that most of the “collisions” happen at the children’s level for a pedigree vertex.

Validating the injection condition is equivalent to solving the well-known Hall’s marriage problem (Hall 1987). Here, we verify this condition directly, which may seem inefficient at first, as the time complexity is exponential, but offers us the benefit of reusable computations (see next section).

#### Exploring child configurations

Taking the Cartesian product for exploring all the children assignment combinations for polytomies is numerically expensive and is, in many cases, not necessary. In this section, we describe a few alternative approaches. The optimal approach depends on the size of the polytomy and the number of candidate assignments per child.

The most straightforward optimization is to eliminate assignments based on a subset of the children. For example, suppose that the chosen vertex *t* has three children: *A* = {*a*_1_, *a*_2_, *a*_3_}. Assume that we have the following assignment space:

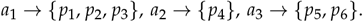

Before exploring every distinct assignment for this triplet, we can solve the same problem for a subset {*a*_2_, *a*_3_} ⊂ *A*. If *p*_4_ and *p*_6_ have no (potential) pedigree MRCAs, we do not need to verify the assignments involving *a*_1_. Having found the “promising” MRCA assignments for *a*_2_ and *a*_3_, we can then proceed by finding those who are also consistent with *a*_1_.

If we intend to use maximum flow for verifying the potential pedigree MRCA condition, we would have to perform it for {*p*_4_, *p*_6_} and every potential pedigree MRCA and then do the same with any third assignment to *a*_1_, which is in some way redundant. With this specific approach, it turns out that the Hall’s condition may offer a significant benefit, allowing us to cache the results for the {*p*_4_, *p*_6_} assignment and reuse it for any further extension of this sub-assignment.

For large polytomies, a general approach would be to explore all the combinations in a DFS manner, caching the intermediate results and trimming all the ‘verification paths’ whenever we find zero valid candidates for *t*.

#### Additional optimization

Finally, to make every single verification faster, we perform additional preprocessing of the graph. For every vertex *v* ∈ *P* and for every its ancestor *a*, we have a map returning the children of *a* through which *v* can reach *a*.

Apart from that, we can also take advantage of the natural symmetry of the problem (see Figure 2) and avoid the whole search completely for one of the ploids of the same individual (in other words, perform all the steps only for one of the ploids).

#### Notation dictionary

**Table 1.** Notation dictionary for symbols used in the text.

| Symbol | Description |
| --- | --- |
| $P$ | Pedigree |
| $T$ | A tree / coalescent tree |
| $L_T$ | Leaf vertices in a graph $T$ |
| $g$ | Initial mapping (i.e., function mapping the leaf vertices in a tree to proband vertices in a pedigree) |
| $\Gamma$ | An edge alignment |
| $f$ | A vertex alignment |
| $\Gamma_f$ | An edge alignment corresponding to a vertex alignment $f$ |
| $T_v$ | A subtree of a tree $T$ where $v$ is the root |
| $\lambda$ | The rate parameter in the Poisson distribution used to generate pedigree errors |
| $D$ | A directed graph |
| $V(D)$ | Vertex set of a directed graph $D$ |
| $E(D)$ | The set of directed edges/arcs of a directed graph $D$ |
| $N^-(u)$ | In-neighborhood of a vertex $u$ |
| $N^+(u)$ | Out-neighborhood of a vertex $u$ |
| $d_D(u)$ | The set of all the descendants of a vertex $u$ in a digraph $D$ |
| $\mathcal{P}_{uv}$ | Set of all directed paths from $u$ to $v$ |
| $\mathcal{P}(D)$ | Set of all directed paths in a digraph $D$ |
| $r(T)$ | Root of a tree $T$ |
| $\pi_T(v)$ | Unique parent of a vertex $v$ in a tree $T$ |
| $c$ | Capacity function |
| $s$ | Source vertex in a flow network |
| $t$ | Target vertex in a flow network |
| $h$ | Flow map |
| $h^-(v)$ | In-flow of a vertex $v$ |
| $h^+(v)$ | Out-flow of a vertex $v$ |
| $v'$ | Access vertex of a vertex $v$ |
| $\alpha$ | Function mapping a vertex $v$ to its non-access version in a flow network graph $H_f$ |

## Supplement

**Figure S1.**
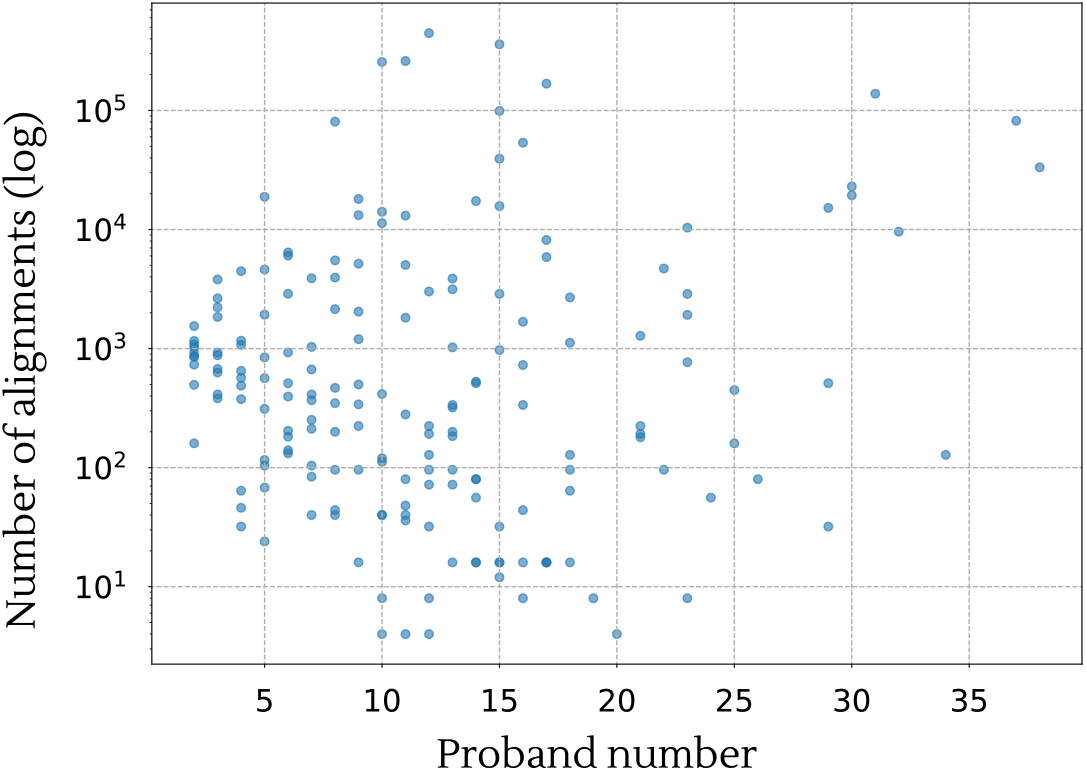
Number of alignments vs proband number (phased regime).

**Figure S2.**
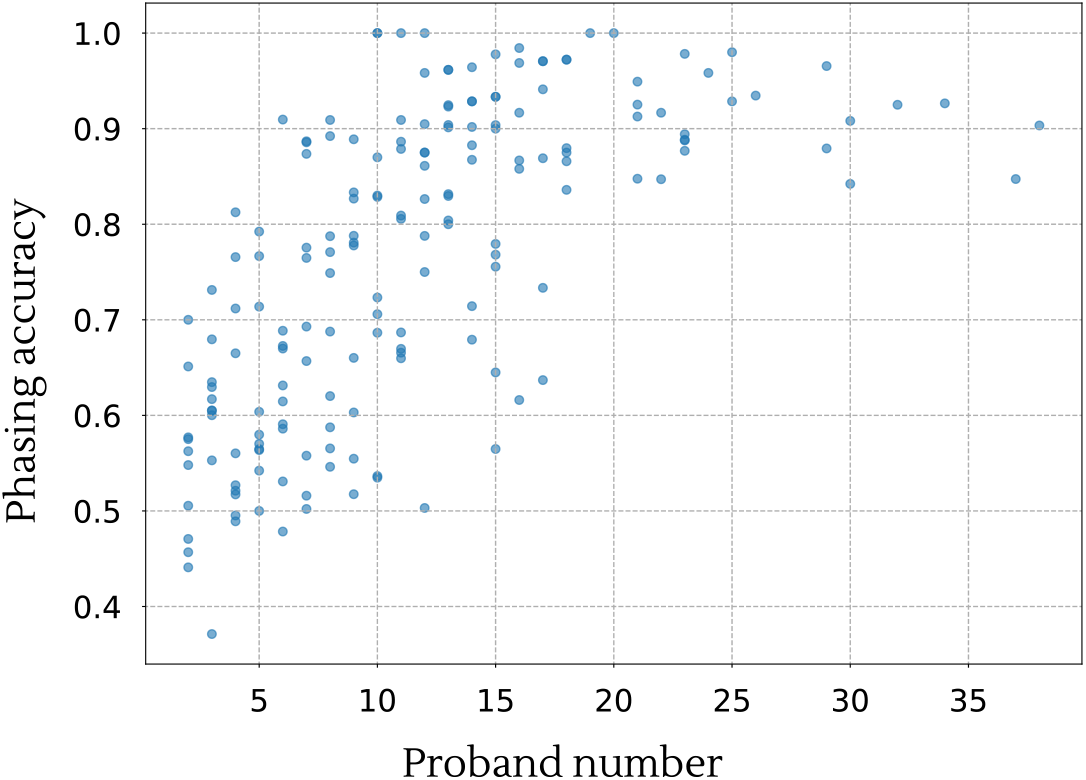
(Unweighted) phasing probability increases with the proband number.

**Figure S3.**
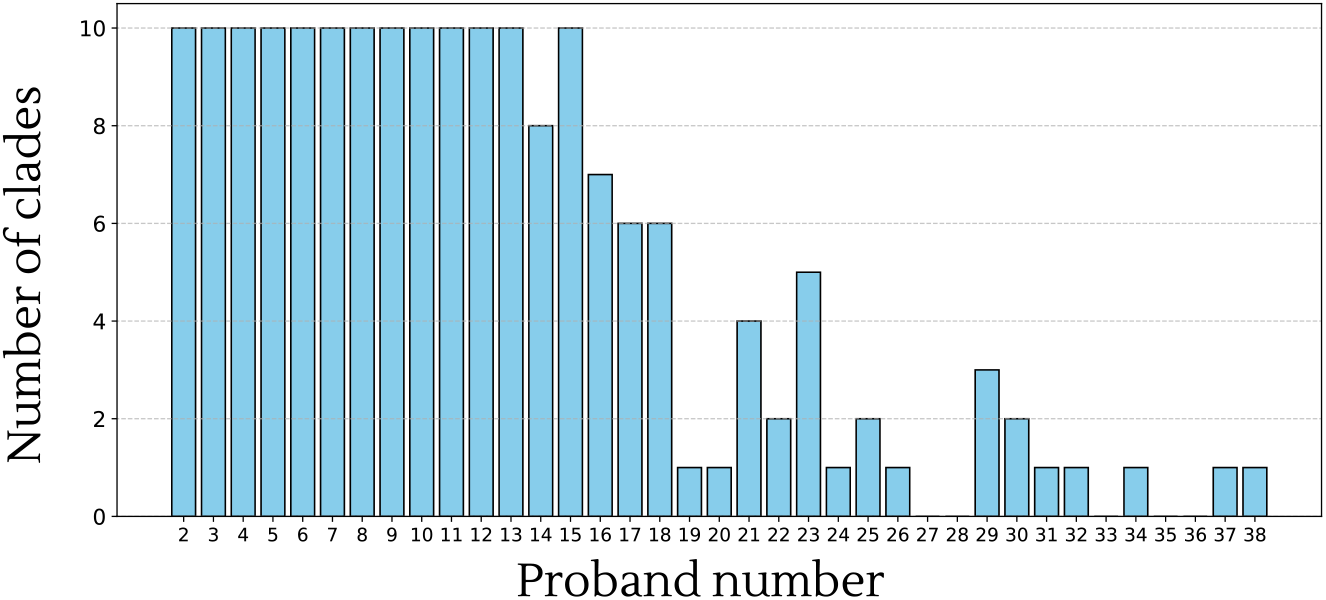
The number of clades simulated in Simulation setup for each proband number.

